# Developing Biosensors for Specific Assessment of *Trans*-translation in *Pseudomonas aeruginosa*

**DOI:** 10.1101/2024.08.30.610505

**Authors:** Bastien L’Hermitte, Thomas Chauvet, Sylvie Georgeault-Daguenet, Nicolas Le Yondre, Philippe Jehan, Reynald Gillet, Christine Baysse

**Affiliations:** Univ. Rennes, CNRS, Institut de Génétique et Développement de Rennes (IGDR) UMR6290, 35000 Rennes, France

## Abstract

*Trans*-translation is a crucial bacterial process and a target for new antibiotics. We developed two *Pseudomonas aeruginosa* biosensor strains that detect *trans*-translation inhibitors by exploiting the bacterium’s natural red fluorescence, linked to protoporphyrin IX accumulation. The first biosensor monitors tmRNA-SmpB-mediated tagging, while the second serves as control for biosensor 1 by keeping track of ClpP1-related proteolysis and porphyrin biosynthesis. Validation through gene deletions and complementation confirmed their specificity. These biosensors were effective in screening antibiotics and designed inhibitors, demonstrating their potential for high-throughput identification of *trans*-translation inhibitors in drug-resistant *P. aeruginosa*.

## 1 Introduction

*Tran*s-translation refers to the molecular process that allows the release of ribosomes which are blocked on mRNAs lacking stop codons. It allows for the degradation of these faulty mRNAs and proteins and promotes canonical recycling of all the other partners (Guyomar et al., 2021). This major rescue pathway has been identified in more than 99% of bacterial genomes and is mediated by transfer-messenger RNA (tmRNA) associated with SmpB protein (Hudson & Williams, 2015). During this process, incomplete nascent peptides are tagged for proteolysis thanks to the messenger-like domain (MLD) of the tmRNA, problematic mRNAs are degraded, and the previously stalled ribosomes are recycled (for a recent review, see D’Urso et al., 2023). In many bacterial species this control machinery is essential for their survival and thus it is an excellent target for the development of antimicrobials compounds (Campos-Silva et al., 2021; Keiler & Feaga, 2014). For species in which this system is not essential the deletion generally diminished the virulence, the tolerance to antibiotics, and more globally, the resistance to stresses (Morita et al., 2006; Ren et al., 2019). It may even prevent the resuscitation of persistent cells as demonstrated for *Escherichia coli* (Yamasaki et al., 2020).

The World Health Organization (WHO) has designated seven “ESKAPEE” pathogenic bacteria (*Enterococcus faecium, Staphylococcus aureus, Klebsiella pneumoniae, Acinetobacter baumannii, Pseudomonas aeruginosa, Enterobacter spp.,* and *Escherichia coli*) as critical targets for drug discovery. Indeed, these highly virulent and resistant bacteria are the leading cause of hospital-acquired infections worldwide. Among these pathogens, the resistance of *P. aeruginosa* to aminoglycosides, quinolones, and β-lactams is of particular concern (Pang et al., 2019). Moreover, due to the emergence of carbapenem-resistant strains, the WHO has identified carbapenem-resistant *P. aeruginosa* as one of three bacterial species for which there is a critical need to develop new antibiotics to treat infections (World Health Organization, 2017, 2024). *P. aeruginosa* possesses the canonical tmRNA/SmpB *trans*-translation system but also genes encoding putative alternative ribosome rescue factors, ArfA and ArfB. Therefore, absence of *trans*-translation does not affect its viability under physiological conditions (Ren et al., 2019) but rather its virulence and adaptation to stressful conditions. Indeed, it has been reported that the *trans*-translation system is crucial for *P. aeruginosa* tolerance to azithromycin and several aminoglycoside antibiotics (Ren et al., 2019). Additionally, the genes encoding tmRNA (*ssrA*) and SmpB showed increased expression following exposure to tobramycin, suggesting a potential role of *trans*-translation for adapting to antibiotics (Cianciulli Sesso et al., 2021).

While *in vitro* high-throughput screening assays to identify *trans-*translation inhibitory molecules in bacteria of the ESKAPEE group have been recently developed (Guyomar et al., 2020; Thépaut et al., 2021), *in vivo* screening systems in living bacterial cells were developed in *E. coli* only (Macé et al., 2017; Ramadoss et al., 2013). These systems are based on replicative plasmids in *E. coli*, encoding either luciferase, mCherry or GFP reporters. These systems require ampicillin for plasmid maintenance and can only be used in bacteria that do not naturally produce green or red fluorescence. Therefore, they are not directly adaptable to *P. aeruginosa* because of its natural resistance to many antibiotics and because this bacterium naturally produces green fluorescence, particularly pronounced in iron-deficient environments, due to the synthesis of the siderophores pyoverdine and pyochelin (Cornelis & Dingemans, 2013). *P. aeruginosa* can also produce red fluorescence related to the accumulation of porphyrin derivatives, detectable in a wild-type strain on medium supplemented with high concentrations of **γ**-aminolevulinic acid, a precursor for the heme biosynthetic pathway (Jones et al., 2020). Protoporphyrin IX is the direct precursor of heme (non-fluorescent) and transformation depends on the activity of the ferrochelatase encoded by the *hemH* gene (also known as *ppfC*). It has been demonstrated that a *hemH* mutant of *Pseudomonas fluorescens* produces a red fluorescence easily detectable on colony and measurable in lysate at an excitation of 405 nm, and an emission at 630 nm (Baysse et al., 2001).

Therefore, in this study, we used this ability of *P. aeruginosa* to naturally produce this endogenous red fluorescent pigment to construct *P. aeruginosa* biosensor strains capable of monitoring *trans*-translation, without the need for fusion with a fluorophore on a plasmid requiring antibiotic selection pressure for its maintenance.

## 2 Material and Methods

### 2.1 Strains, Plasmids, Primers, and Growth Conditions

All the strains, plasmids and primers are listed in table 1. *E. coli* strains used for plasmid construction have been cultivated in Luria-Bertani broth (LB) and LB agar medium containing either gentamycin (10µg/mL), ampicillin (100µg/mL) or chloramphenicol (20µg/mL). *P. aeruginosa* strains have been grown in LB, casamino acids medium (CAA) (casein peptone 5g/L, K_2_HPO_4_ 0.9g/L, MgSO_4_•7H_2_O 0.25g/L), CAA agar, supplemented with sucrose (15% w/v) if necessary and CN agar for *Pseudomonas* (Grosseron). Gentamycin (100µg/mL), carbenicillin (300µg/mL) or chloramphenicol (100µg/mL) were added if needed. For solid media, 1.5% (p/v) agar was used.

**Table 1:**
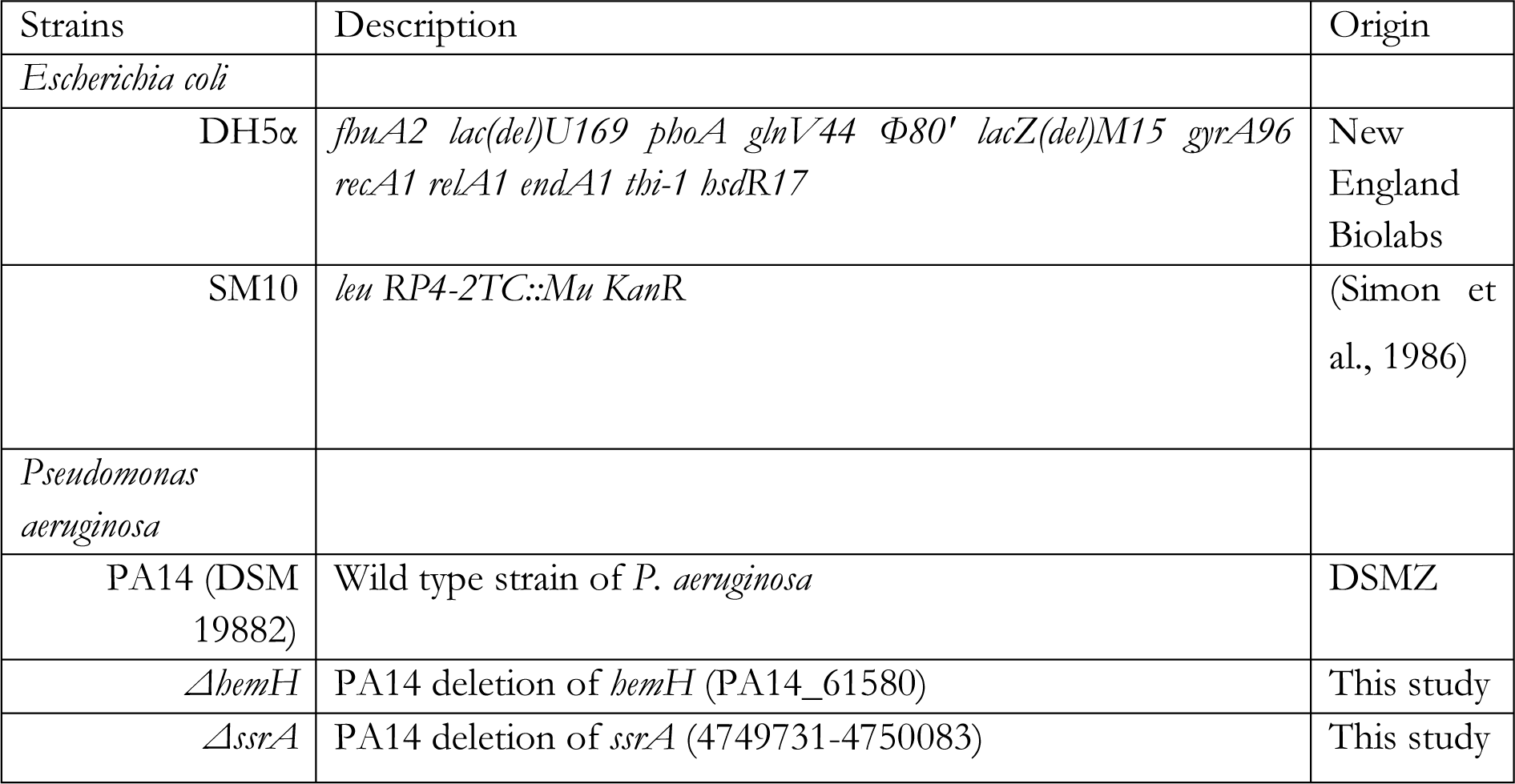
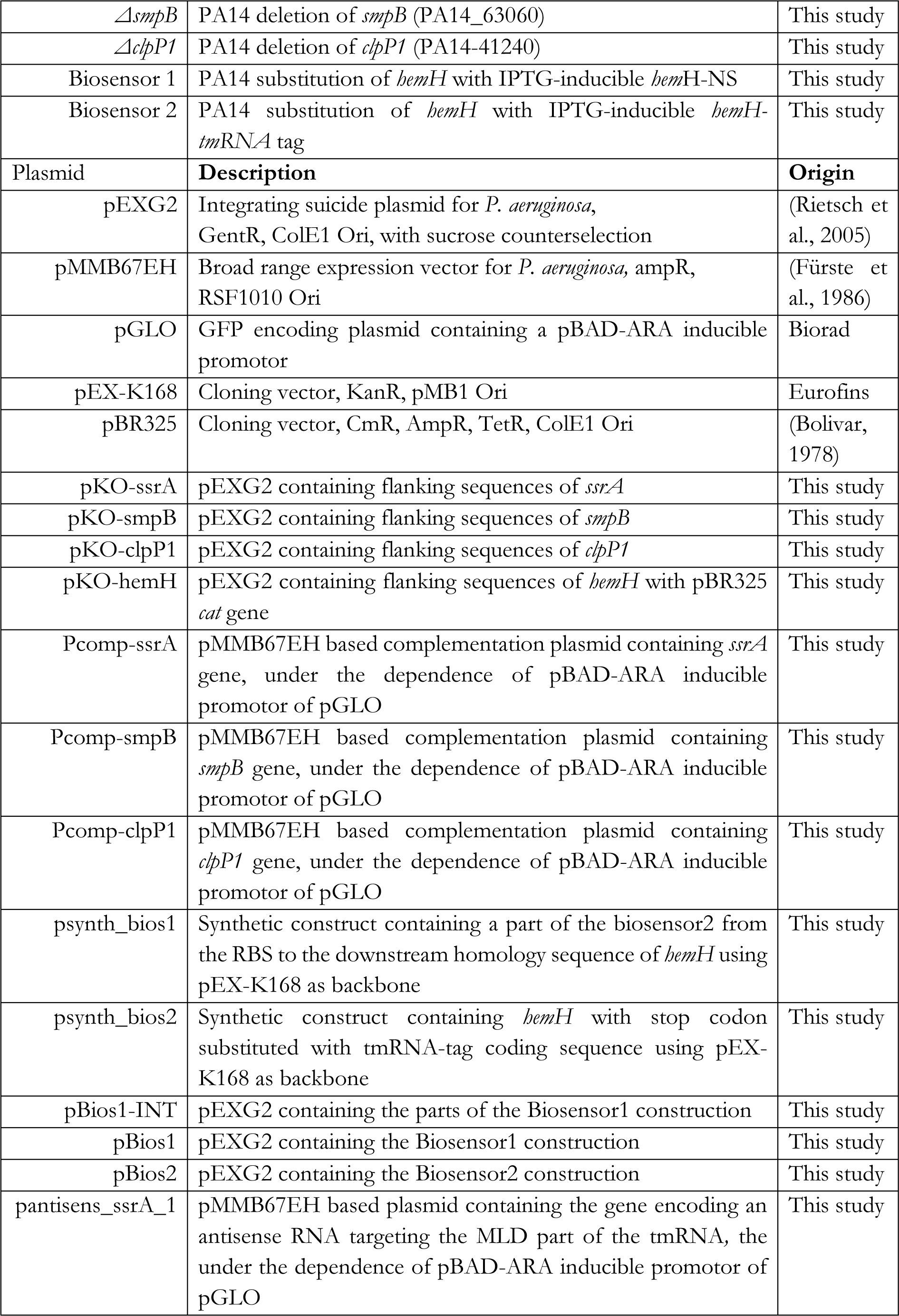
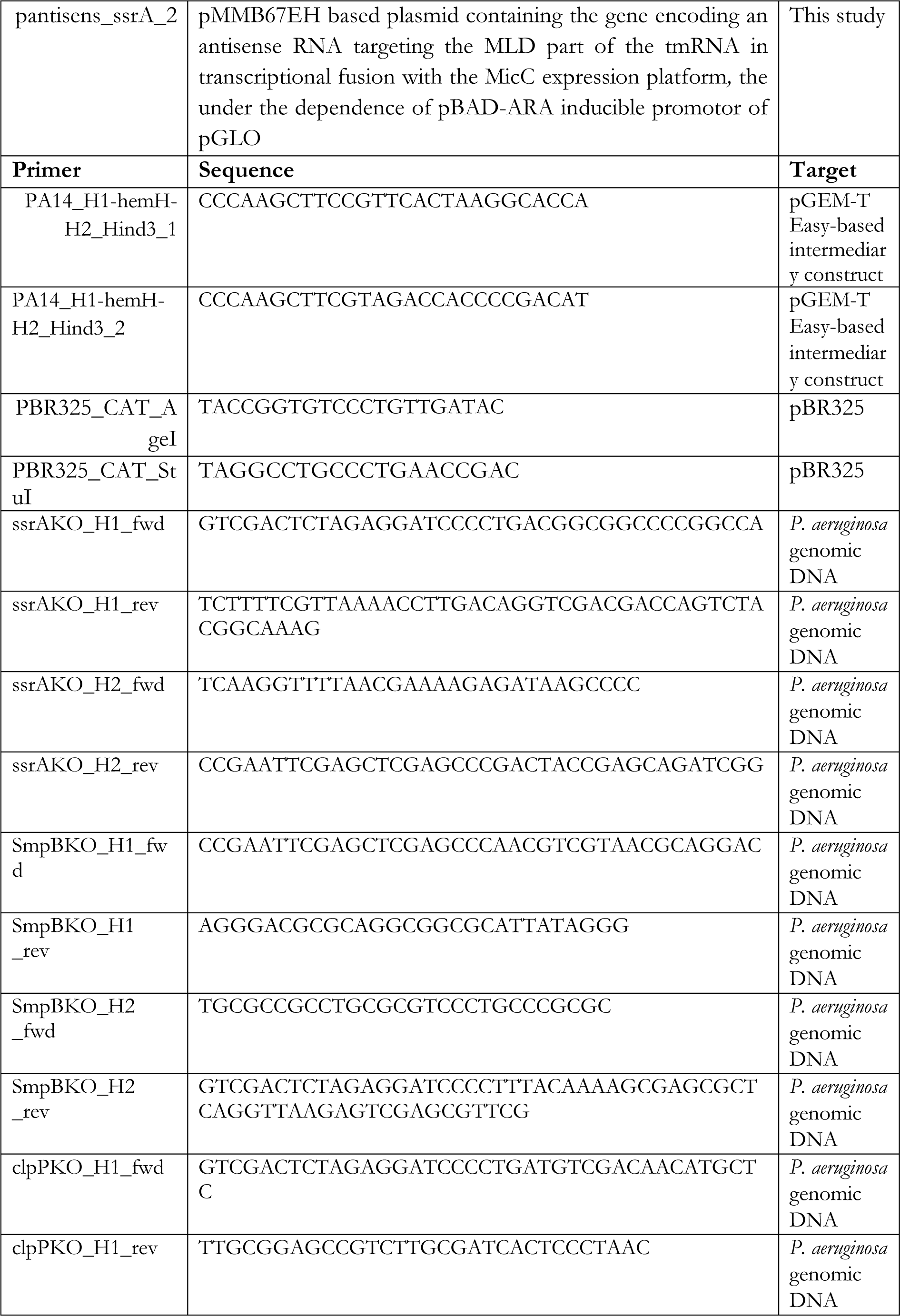
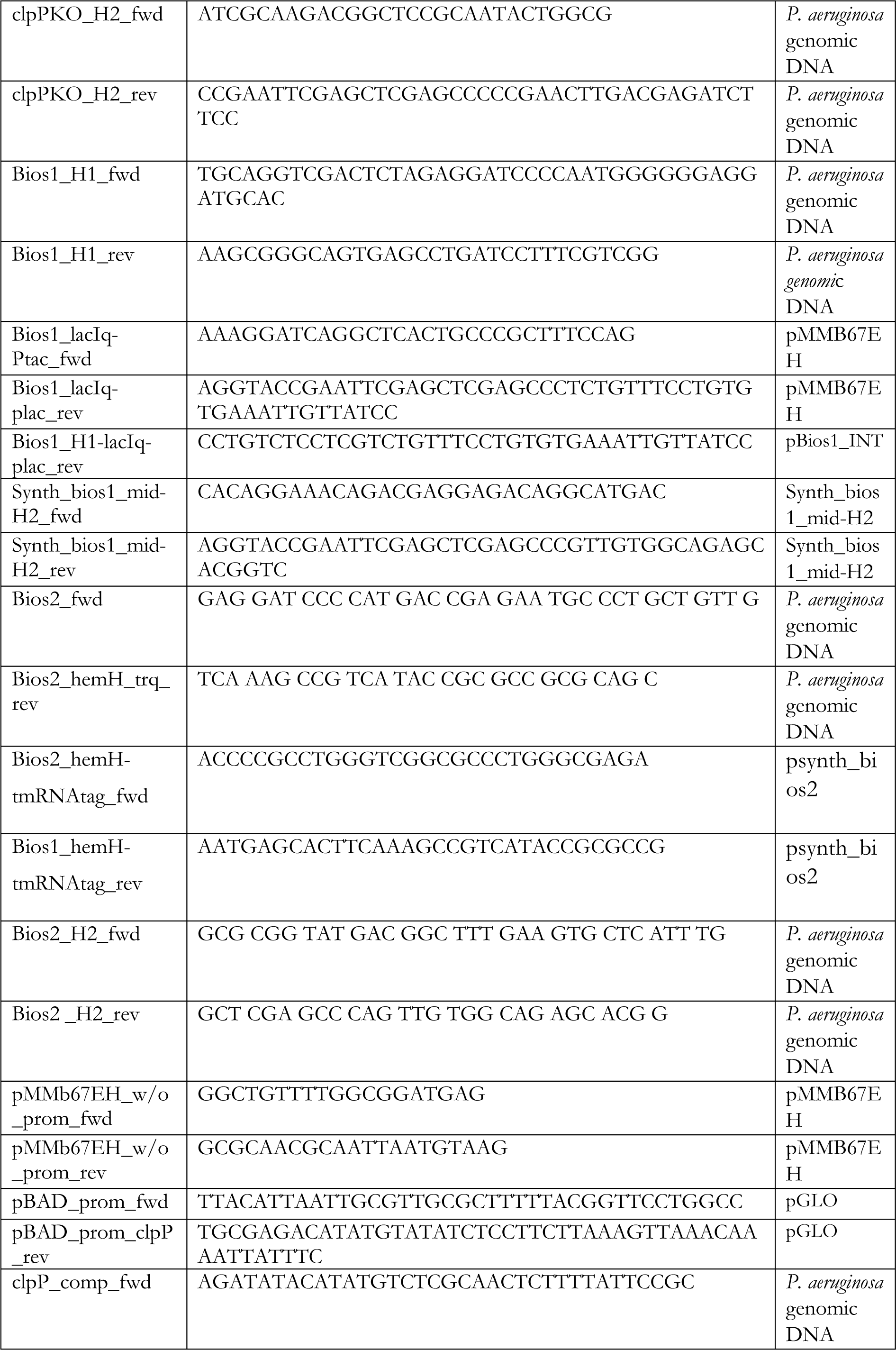
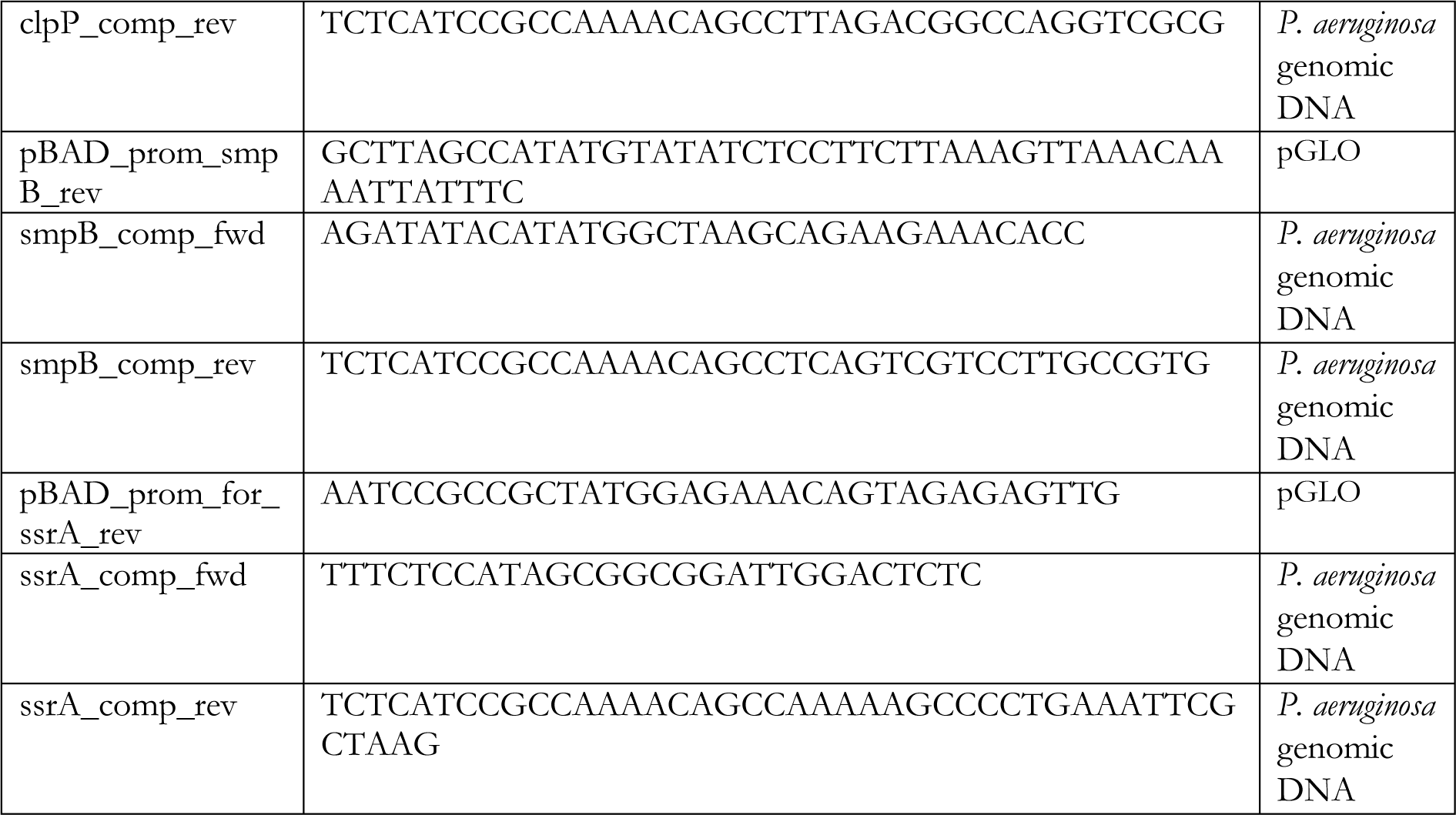
strains, plasmids and primers.

Biosensor testing medium (Btm) is composed of CAA supplemented with 40µM of FeSO_4_, 0.5mM aminolevulinic acid, 10µM isopropyl β-D-1-thiogalactopyranoside (IPTG) and, for complementation assays, 10mM arabinose.

### 2.2 Plasmids and strains construction (supplementary Fig.S1)

To assess the viability of the *P. aeruginosa hemH* mutant and its capacity to produce red fluorescence, the central part of the gene was replaced with the gene coding for chloramphenicol acetyltransferase (CAT) from the pBR325 plasmid, to facilitate selection. The H1-*hemH*-H2 fragment was amplified from *P. aeruginosa* PA14 genomic DNA using the PA14_H1-*hemH*-H2_*Hind*III_1 and PA14_H1-*hemH*-H2_*Hind*III_2 primers, then cloned via TA cloning into the pGEM-T Easy system. Next, the *cat* cassette, amplified from pBR325 with the PBR325_CAT_AgeI and PBR325_CAT_*Stu*I primers, was cloned into the pGEM-T Easy H1-*hemH*-H2 at the *Age*I and *Stu*I restriction sites. Finally, the H1-*hemH*::*cat*-H2 *Hind*III fragment was cloned into the pEXG2 plasmid at the *Hind*III site, creating pKO-hemH. All restriction enzymes used for this construction were purchased from New England Biolabs.

The pKO plasmid series, except for pKO-hemH, was constructed by cloning fragments flanking the gene to be deleted into the SmaI site of pEXG2. The fragments were obtained by PCR using Q5® DNA polymerase (New England Biolabs) and the primers listed in Table 1. The amplification products were purified using the Monarch® PCR & DNA Cleanup Kit (New England Biolabs), and the final constructs were obtained using HiFi assembly (New England Biolabs) and the NEBuilder® assembly tool, according to the manufacturer’s instructions.

The pBios1-INT plasmid, which uses pEXG2 as a backbone, contains the upstream homology sequence of *hemH* (H1), obtained using the Bios1_H1_fwd and Bios1_H1_rev primers with genomic DNA as the template. Downstream of this sequence, the plasmid includes the IPTG-inducible *lacI*q-Ptac-*lac*O promoter, derived from the pMMB67EH plasmid (Fürste et al., 1986), and amplified using the Bios1_lacIq-Ptac_fwd and Bios1_lacIq-plac_rev primers. From pBios1-INT, the H1-*lacI*q-Ptac*-lac*O fragment was amplified using the Bios1_H1_fwd and Bios1_H1-lacIq-plac_rev primers and cloned into pEXG2, upstream of the synthetic fragment present on psynth_bios2_mid, amplified using the Synth_bios1_mid-H2_fwd and Synth_bios1_mid-H2_rev primers, creating the pBios1 plasmid (Table 1, Fig. S1).

The pBios2 plasmid was obtained by cloning most of the *hemH* gene, amplified using Bios2_fwd and Bios2_hemH_trq_rev, followed by a fragment containing the rest of the *hemH* gene fused with the tmRNA degradation tag, amplified from psynth_bios2 using Bios2_hemH-tmRNAtag_fwd and Bios2_hemH-tmRNAtag_rev primers, and the downstream homology sequence of *hemH* (H2) amplified using the Bios2_H2_fwd and Bios2_H2_rev primers. The pBios2 uses the pEXG2 linearized by *Sma*I as backbone. (Fig. S2)

The pcomp plasmid series has been created using the same supplies as the pKO series. The lactose-inducible promotor and multiple cloning site of the pMMB67EH has been replaced by the arabinose inducible promotor and AraC regulator of the pGLO plasmid. The RBS upstream of the promoter has been conserved if necessary. For each complementation, the corresponding native gene has been cloned downstream of this new promoter.

All mutant strains were obtained by following the method described in Hmelo et al. (2015) using the “puddle” variation and by replacing LBNS medium by CAA.

The inhibitor plasmids pantisens_ssrA_1 and pantisens_ssrA_2, have been created by cloning both the synthetic fragments (Table S1), (Genscript), between *Mlu*I and *Pst*I sites of the pMMB67EH using Hi-T4 DNA Ligase (NEB).

### 2.3 High throughput compound testing procedure

For biosensor testing, *P. aeruginosa* strains were inoculated at an initial OD_600_ of 0.05 in testing medium Btm. Tests were performed in 96 well flat bottom plates (ref: 353072, Falcon). The plate was incubated in the BioTek Synergy HTX Multimode Reader (Agilent) at 37°C. Both the OD_600_ and the red fluorescence intensity (EX: 400nm/ EM: 620nm) were measured.

Data analysis is performed as follows: the OD_600_ is linearized using the formula log_2_(100 * OD_600_). The normalized fluorescence intensity is calculated by dividing the fluorescence intensity (EX: 400nm / EM: 620nm) by the linearized OD_600_. The inflection point of each curve is then determined using the “gcplyr” R package to rephase the curves (Blazanin, 2023). Mean and standard deviation calculations are performed after linearization, normalization, and rephasing. Statistical analysis was performed using a Kruskal-Wallis test followed by a Dunn post-hoc test on the points between 150-350 minutes for Biosensor 1 and between 150-520 minutes for biosensor 2.

For specificity testing of the biosensors, tobramycin, netilmicin and ciprofloxacin have been used at a concentration of 0.083µg/mL, 0.17µg/mL and 0.02µg/mL, respectively.

For fluorescence intensity measurements on the lysate, the cell pellet was disrupted using a mixture of 0.1 M NH_4_OH and acetone in a 9:1 (v/v) ratio.

### 2.4 Mass spectrometry analysis

96 wells plates were inoculated in the previously stated conditions with either wild type or biosensor2 *P. aeruginosa* strains. After 250 minutes of incubation, the cells were collected by centrifugation and subsequently lysed by vortexing in 6M formic acid. The samples were freeze dried and then resuspended in either acetone plus 1% (v/v) formic acid or DMSO. The HPLC system used was an UltiMate 3000 HPLC RSLC cap system coupled to a Q Exactive mass spectrometer, equipped with a HESI II source (Thermo Fisher Scientific, Bremen, Germany). Samples were acquired either in direct infusion mass spectrometry (DI-MS) at 5 µl/min with a syringe pump or liquid chromatography coupled with mass spectrometry (LC-MS), after loading on a trap column of 1 µl. Mass spectra (MS1) or tandem mass spectra (MS2) were recorded at a resolution of 140,000 at m/z 200.

## 3 Results

### 3.1 Red-fluorescence of *P. aeruginosa ΔhemH*

To develop a cell-based system for monitoring *trans*-translation, we leveraged the natural red fluorescence of the *P. aeruginosa ΔhemH* mutant, which arises due to the accumulation of protoporphyrin IX (PPIX), the substrate of ferrochelatase encoded by the *hemH* gene. To enhance PPIX production and bypass the regulation of the PPIX biosynthesis pathway, which primarily occurs at the *hemA* gene encoding the glutamyl-tRNA reductase, we supplemented the growth medium with 5-aminolevulinic acid (ALA), a precursor in the tetrapyrrole biosynthetic pathway, which is upstream of PPIX and downstream of glutamate1 semialdehyde, the product of glutamyl-tRNA reductase (Fig. S1) (Choby & Skaar, 2016; Hungerer et al., 1995). The fluorescence intensity of the Δ*hemH* mutant was observed to be directly correlated with the concentration of ALA in the medium, confirming the role of ALA in modulating the red fluorescence signal (FigS1). Development of two biosensor strains to evaluate *trans*-translation inhibition in *P. aeruginosa*.

To investigate the *trans*-translation process in *P. aeruginosa*, we engineered two distinct biosensor strains. The first strain, named biosensor 1, was designed to assess the effect of molecules on the tagging efficiency of *trans*-translation mediated by the SmpB-tmRNA system, while the second strain, biosensor 2, serves as a control to evaluate either the effect on the degradation of tmRNA-tagged proteins by the ClpP1-containing protease complex or the effect on PPIX synthesis. Both strains are based on modifications of the *hemH* gene, which encodes the ferrochelatase, which catalyzes the insertion of ferrous iron into protoporphyrin IX.

### 3.2 Biosensor 1: a tool to measure *trans*-translation activity

To specifically induce red fluorescence under conditions where *trans*-translation is not inhibited, we modified the *hemH* gene to produce a protein that becomes a substrate for tmRNA-SmpB-mediated tagging. The native *hemH* gene was replaced by a modified version (*hemH*-ns) lacking its translation termination codon (Fig.S3). This modified gene was placed under the control of the IPTG-inducible *lacI*q-Ptac-*lac*O promoter derived from the pMMB67EH vector (Fürste et al., 1986), with two Rho-independent 5S ribosomal RNA (PA14_55620, Winsor et al., 2016) transcription termination sequences ensuring proper transcriptional termination (see material and methods). The resulting stalled ribosome complex at the end of the *hemH*-ns mRNA requires resolution by *trans*-translation. If *trans*-translation is functional, the tagged ferrochelatase is addressed for degradation by the ClpP1-containing proteases (Gottesman et al., 1998), leading to the accumulation of protoporphyrin IX (PPIX) and the emission of a red fluorescent signal in the Biosensor 1 strain (Fig. 1).

**Figure 1:**
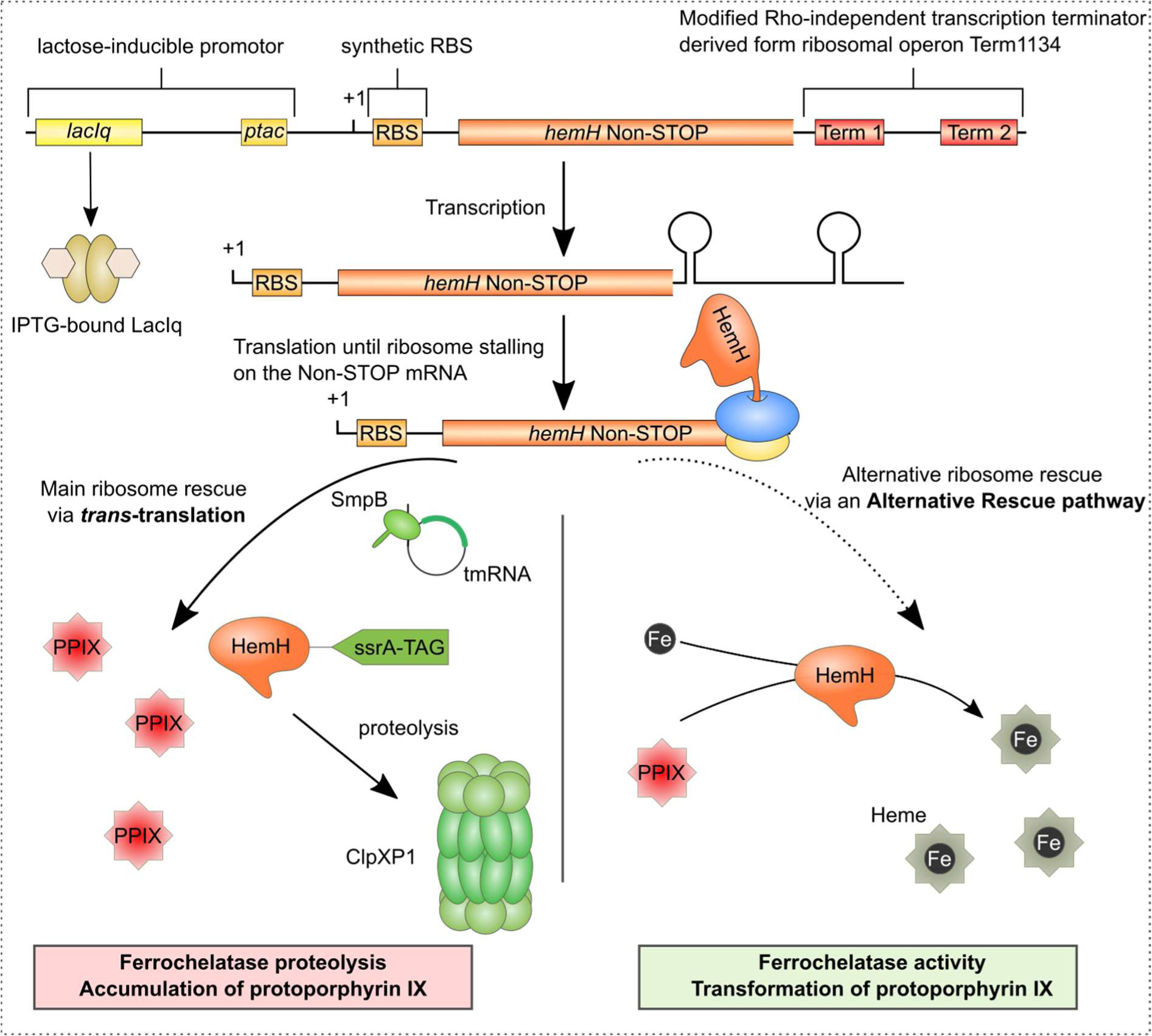
Schematic Representation of Biosensor 1 Functionality. Upon induction, the synthetic construct generates a non-stop mRNA, which necessitates resolution by *trans*-translation. This process tags the newly synthesized HemH protein for degradation by adding the tm-RNA-TAG (“ssrA-TAG”) at the C-terminus. Degradation of ferrochelatase leads to the accumulation of protoporphyrin IX and results in red fluorescence within the bacterial cell. If *trans*-translation is absent, inhibited, or overwhelmed, the HemH protein remains untagged and free, enabling the conversion of protoporphyrin IX into heme, thus reducing the red fluorescence.

First, it was important to optimize the culture conditions for biosensor 1. We tested the concentrations of three key compounds: iron (a substrate for ferrochelatase in the production of heme from PPIX), aminolevulinate (a precursor in the heme biosynthetic pathway), and IPTG (an inducer of the *hemH*-ns gene in biosensor 1). CAA medium was chosen as the base medium to effectively control iron concentration. To evaluate the performance of each tested combination, we measured the difference in fluorescence signal at the 250-minute mark between biosensor 1 and its Δ*ssrA* mutant strain. Ideally, we aimed for a high fluorescence signal in biosensor 1 and a significant difference compared to its Δ*ssrA* mutant strain. Based on the results, the optimal combination was identified as follows: 10 µM IPTG, 0.5 mM aminolevulinate, and 40 µM FeSO₄ (Fig. 2).

**Figure 2:**
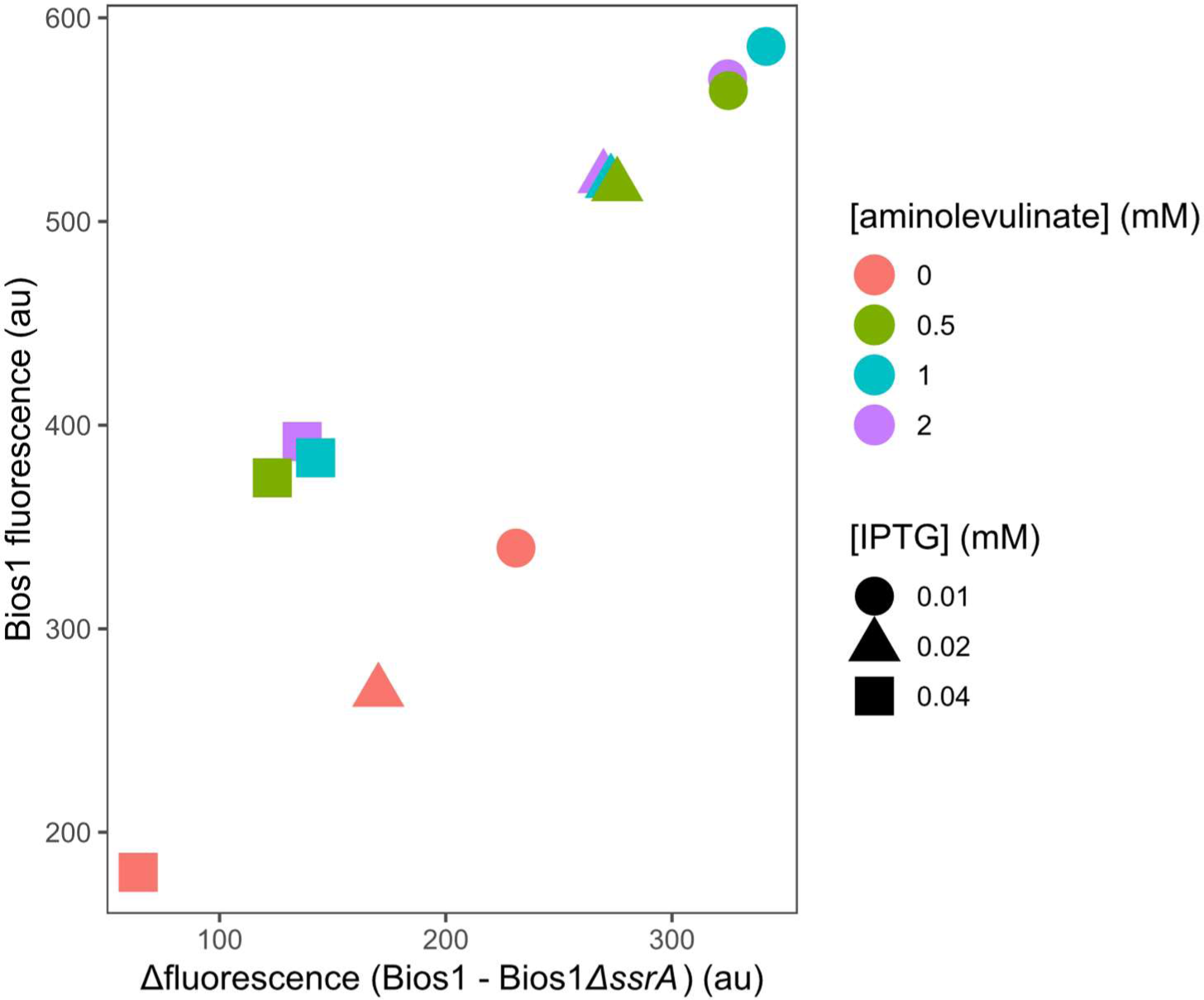
Fluorescence level differences between Biosensor 1 and its ΔssrA mutant strains at the 250-minute mark, illustrating the maximum yet stable signal under the tested growth conditions. Data is shown only for [FeSO₄] = 40 µM, as this parameter did not influence fluorescence intensity.

To validate the correlation between fluorescence and *trans*-translation, we constructed biosensor 1 *ΔsmpB* and biosensor 1 *ΔssrA* strains, for which *trans*-translation is disrupted. In the wild-type (WT) background, tmRNA and SmpB are active, leading to PPIX accumulation. In contrast, in the *ΔsmpB* or *ΔssrA* mutant strains, a reduction in red fluorescence is anticipated because the untagged HemH is not targeted for degradation, allowing PPIX conversion to heme (Fig. 1). To ensure that the observed decrease in red fluorescence in the mutated biosensor 1 strains was solely due to the deletion of *ssrA* or *smpB*, we performed complementation experiments using specially designed plasmids carrying the missing gene under the control of an arabinose-inducible promoter. Complementation of the biosensor 1 *ΔssrA* strain fully restored the biosensor 1 phenotype, while complementation of the biosensor 1 *ΔsmpB* strain only partially restored it. This partial restoration is likely due to the reduced growth rate caused by the metabolic burden of the pcomp-smpB plasmid, as observed in both the biosensor 1 and biosensor 1 *ΔsmpB* complementation strains (Fig. 3).

**Figure 3:**
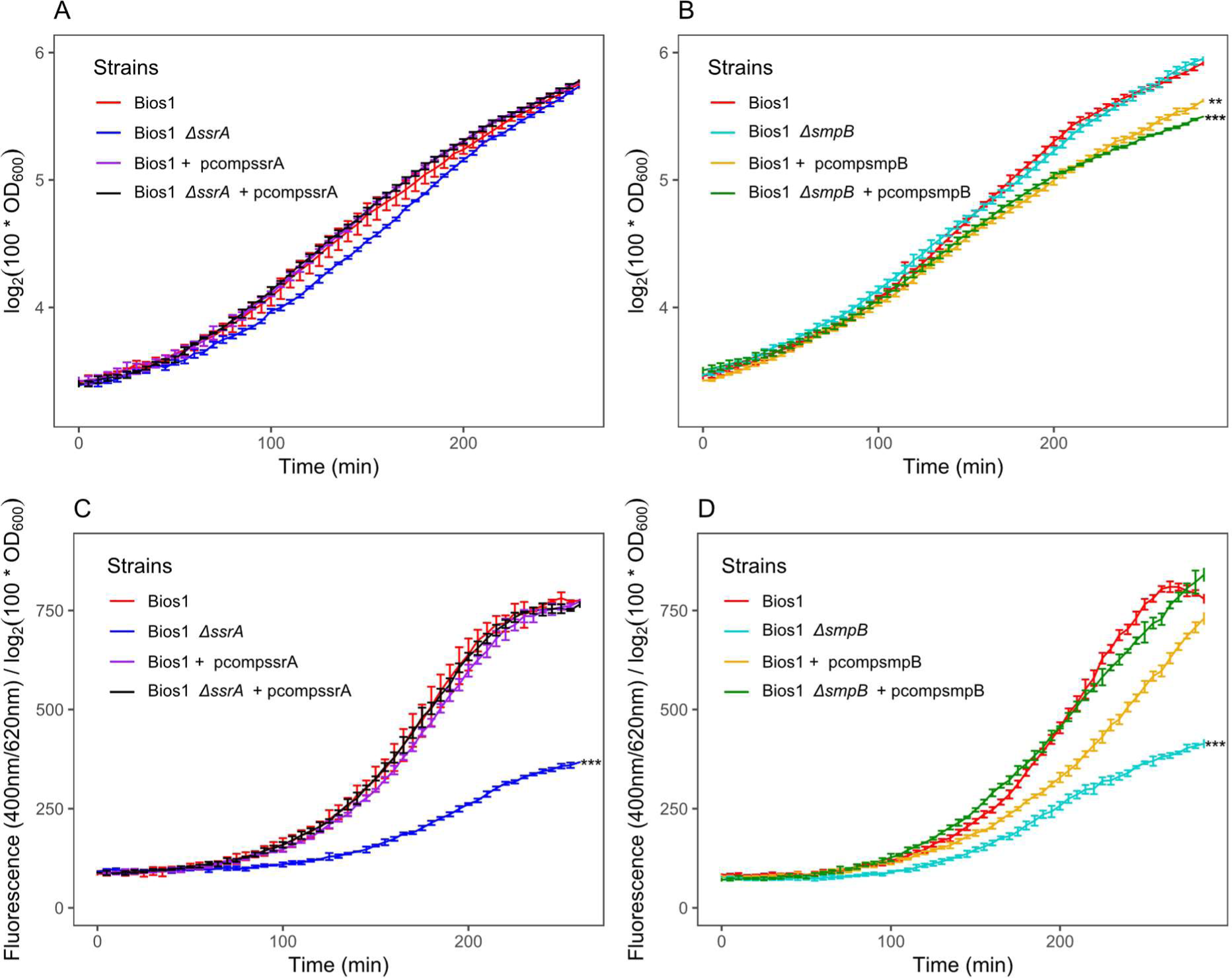
Validation of the biosensor1. (A-B) Growth curve (optical density at 600nm according to time in minutes) of biosensor 1, its *ssrA* (A) or *smpB* (B) mutant strain and the corresponding complemented strains. (C-D) Fluorescence normalized by the optical density according to time in minutes of biosensor1, its *ssrA* (C) or *smpB* (D) mutant strain and the corresponding complemented strains. Three repeats have been used for mean calculation and standard deviation. (**) p-value <0.01; (***) p-value <0.001 (Kruskal-Wallis test followed with Dunn test).

The conversion of ALA to PPIX is a multistep process that involves the synthesis of various porphyrins, which are organic compounds known for their ability to emit red fluorescence under specific conditions. In *P. aeruginosa*, the heme biosynthetic pathway can diverge to produce several porphyrin derivatives, including PPIX and coproporphyrin III, all of which can contribute to red fluorescence (Huang et al., 2010). To ensure that the red fluorescence observed in biosensor 1 is predominantly due to the accumulation of PPIX, we conducted mass spectrometry analyses. Both direct infusion mass spectrometry (DI-MS) and liquid chromatography-mass spectrometry (LC-MS) were employed to detect and quantify relatively the porphyrins present in both the wild-type and biosensor 1 *P. aeruginosa* strains. Our analyses confirmed the presence of PPIX and coproporphyrins (Fig. 4).

**Figure 4:**
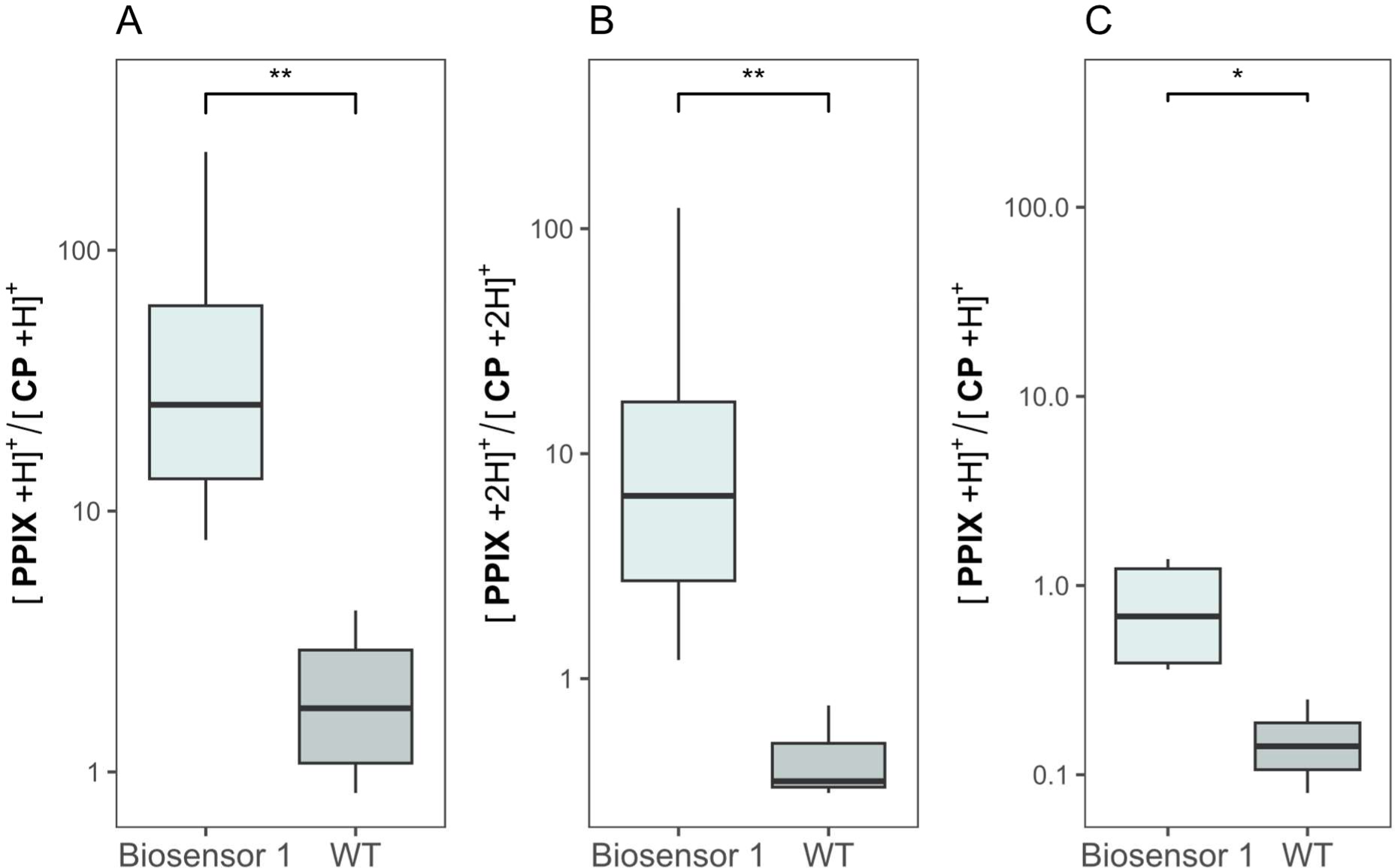
Ratio of protoporphyrin IX and coproporphyrin extracted from the Biosensor 1 and the wild-type (WT). A) ratio of the [X+H] ^+^ species with DMSO as solvent. B) ratio of the [X+2H]^+^ species with DMSO as solvent. C) ratio of the [X+H]^+^species with acetone +1% formic acid as solvent. A) and B) were obtained by DI-MS and C) by LC-MS. (**) p-value <0.01; (*) p-value <0.06 (Kruskal-Wallis test followed with Dunn test).

The ratio of PPIX to coproporphyrins observed in these analyses supports the hypothesis that protoporphyrin IX accumulates in the *P. aeruginosa* biosensor 1 strain, contributing significantly to the red fluorescence detected (Fig. 4). This accumulation is consistent with the conversion of aminolevulinate mainly in PPIX and the role of HemH in converting PPIX to heme.

### 3.3 Biosensor 2: a tool to measure proteolytic activity of ClpP1-containing protease

Biosensor 2 was developed as a control to verify that the molecules inhibiting biosensor 1 fluorescence do not interfere with ClpP1-mediated proteolysis of tmRNA-tagged proteins or PPIX production by disrupting either this biosynthesis pathway or canonical translation rather than *trans*-translation. *P. aeruginosa* possesses two isoforms of the ClpP peptidase: ClpP1 and ClpP2. The *clpP*1 gene is expressed throughout the growth phases, while the *clpP*2 gene is regulated by quorum-sensing signaling molecules and the transcription factor LasR. Their respective roles in *trans*-translation are not well known. However, ClpP1 is an essential component of ClpXP enzymes, which recognize and degrade tmRNA-tagged translation products. For ClpP2 to function as a protease, it must co-assemble with ClpP1 to form a heterocomplex and only the ClpP1 ring of this complex can bind ClpX (Mawla et al., 2021).

To construct biosensor 2, the native *hemH* gene in *P. aeruginosa* PA14 was replaced with the *hemH* gene in translational fusion with the tmRNA-tag of the WT strain (ANDDNYALAA*). When proteolysis mediated by the ClpP1-containing complex is effective, HemH fused with the tmRNA-degradation tag is degraded, leading to the accumulation of PPIX in the cell, resulting in red fluorescence (Fig. 5). As a control to link the fluorescence with proteolytic activity, the *clpP1* gene was deleted in biosensor 2. The *ΔclpP1* mutant does not exhibit an increase in fluorescence over time, unlike biosensor 2 in the WT background, consistent with the previously established role of ClpP1 in degrading GFP-tmRNA-tagged proteins, even in the presence of ClpP2 (Mawla et al., 2021). Further complementation assays confirmed that reintroducing a *trans*-encoded *clpP1* partially restored the wild-type phenotype, although with limited efficiency (Fig. 6). The reduced growth observed during complementation suggests that overexpression of the pleiotropic ClpP1 protease impacts bacterial fitness (data not shown). This validation step confirmed that biosensor 2, which contains the ClpP1 peptidase containing-complex, produces red fluorescence due to the degradation of the HemH-tmRNA-tag fusion protein. The fluorescence decreases significantly in biosensor 2 Δ*clpP1* and should be similarly affected by exogenous inhibition of ClpP1-containing protease or PPIX synthesis.

**Figure 5:**
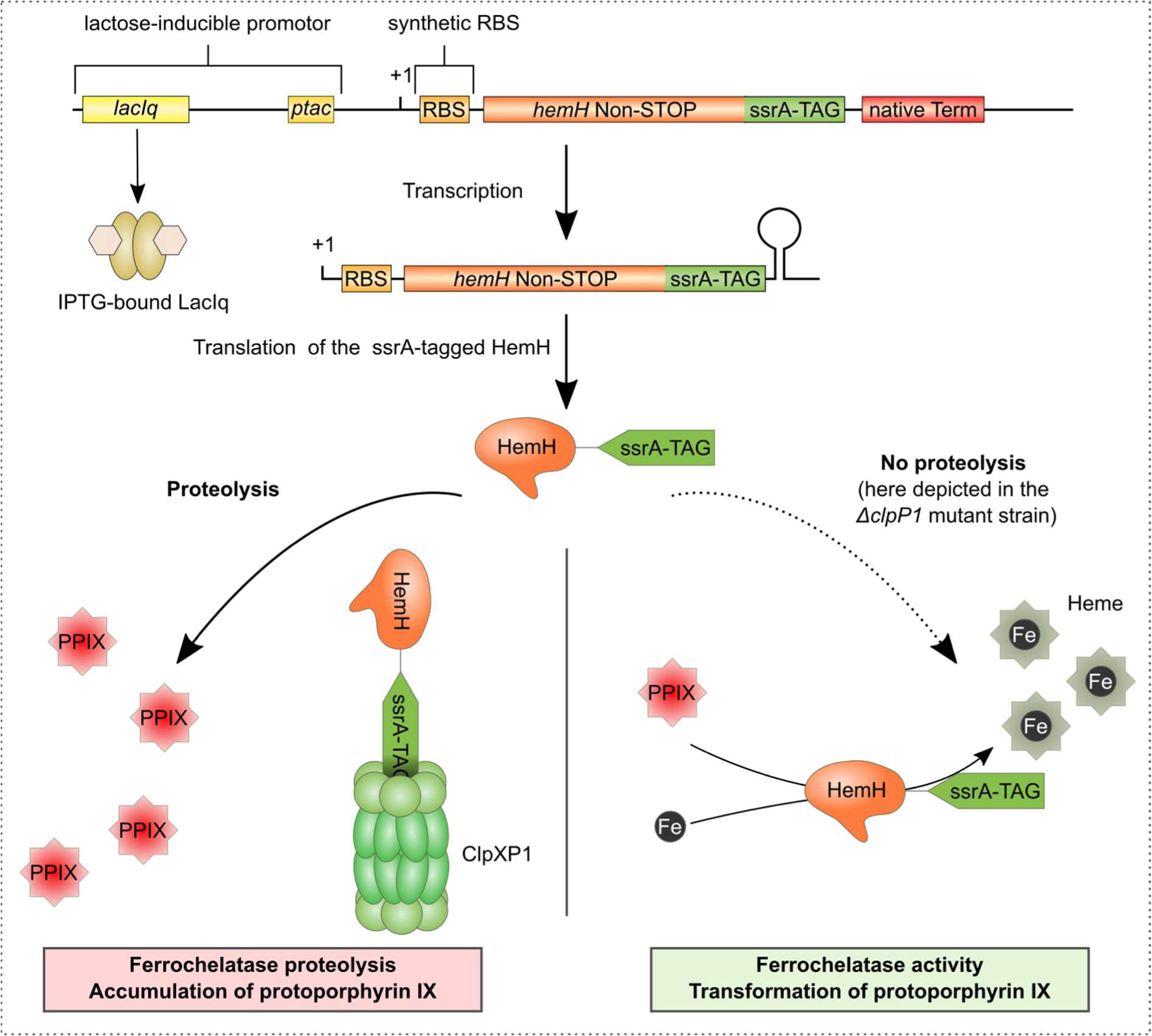
Schematic representation of the functioning of biosensor 2. Upon induction with IPTG, the synthetic construct expresses HemH fused with the tmRNA tag for degradation. This tagged HemH is addressed for degradation to the ClpP1-associated protease complex, leading to the accumulation of protoporphyrin IX, which causes the cell to become red fluorescent. If ClpP1-mediated proteolysis is inhibited or absent, the tagged HemH persists, allowing the conversion of protoporphyrin IX to heme, thereby reducing the red fluorescence. Furthermore, molecules that inhibit the heme biosynthesis pathway will also lead to reduced red fluorescence in biosensor 2.

**Figure 6:**
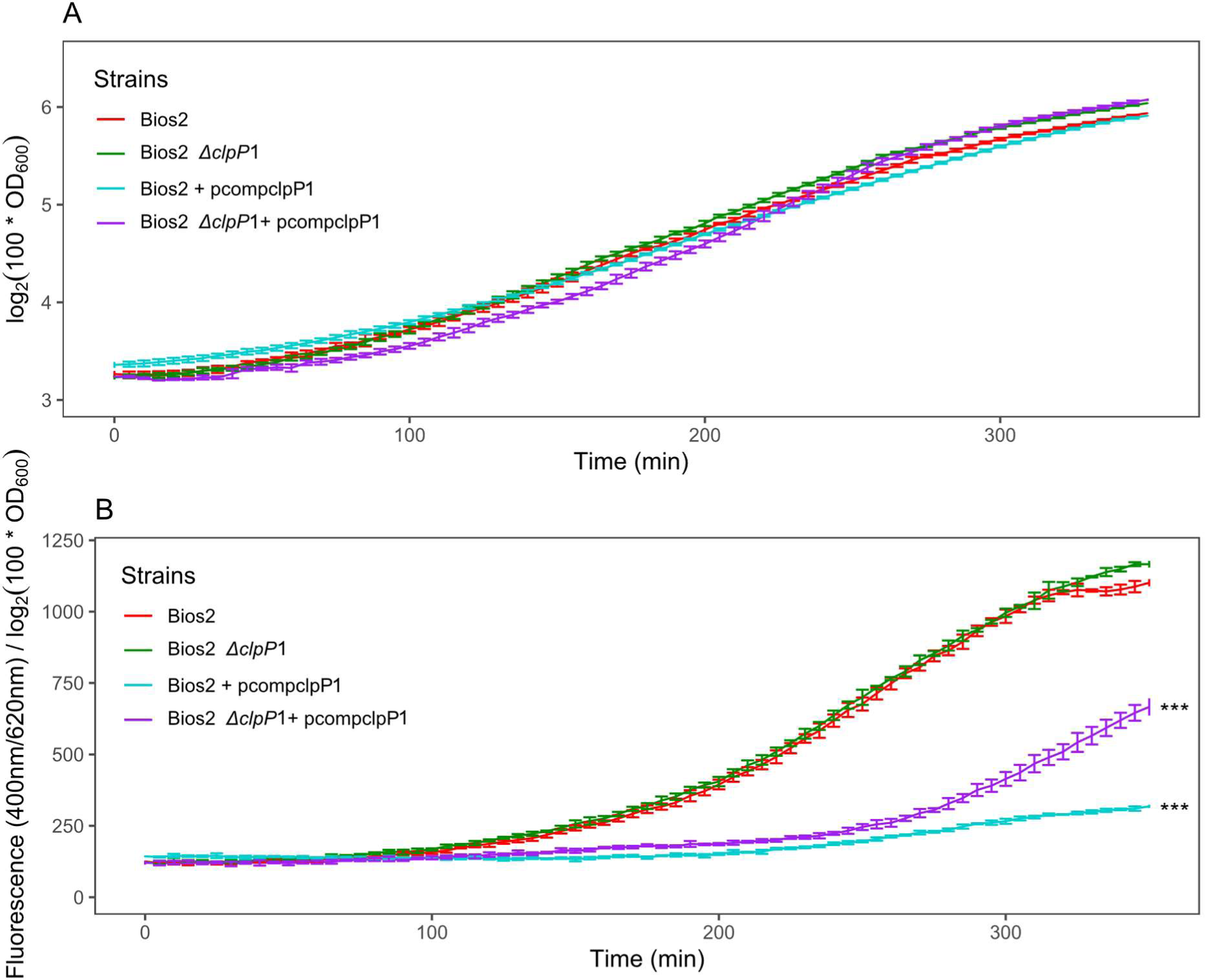
Validation of the biosensor 2. (A) Growth curve (optical density at 600nm according to time in minutes) of biosensor 2, its *clpP1* mutant strain and the corresponding complemented strains. (B) Fluorescence normalized by the optical density according to time in minutes of biosensor 2, its *ΔclpP1* mutant strain and the corresponding complemented strains. Three repeats have been used for mean calculation and standard deviation. (***) p-value <0.001. (Kruskal-Wallis test followed with Dunn test)

### 3.4 Effect of inhibitors on the system

To evaluate the system’s responses under various conditions, we tested its performance in the presence of commonly used antibiotics and specifically designed inhibitors.

#### 3.4.1 Effect of antibiotics on the performance of the biosensors

To ensure the specificity of the biosensors developed for detecting *trans*-translation inhibitors, it was essential to evaluate their performance in the presence of commonly used antibiotics. This evaluation helps rule out any potential false positives that could arise from antibiotic interference with the fluorescent output of the biosensors. Three antibiotics were selected for testing: netilmicin and tobramycin, both targeting the ribosome, and ciprofloxacin, which targets a non-ribosomal site (DNA gyrase). Each antibiotic was administered at one-third of its minimum inhibitory concentration (MIC) to assess whether sub-inhibitory levels would affect the fluorescence of the biosensors. The results demonstrated that none of the tested antibiotics significantly altered the fluorescence levels in either biosensor. This suggests that the biosensors are robust and maintain their specificity, even in the presence of these antibiotics (data not shown).

#### 3.4.2 Effect of specifically designed inhibitors

The impact of specifically designed *trans*-translation inhibitors on the biosensors was investigated with the aim of testing the sensitivity of our detection system. As no molecules specifically targeting *P. aeruginosa trans*-translation have been formally identified yet, we designed two types of inhibitors: antisense RNAs and a peptide derived from the C-terminal tail of SmpB, both already shown to impact *trans*-translation in *E. coli* (Macé et al., 2017). Antisense RNAs, encoded by plasmids pAntisens_1 and pAntisens_2, targeted the mRNA-like domain (MLD) of the tmRNA to inhibit its function in *trans*-translation. The pAntisens_2 construct, enhanced with an *E. coli* MicC sRNA scaffold, was designed to improve the HFQ-mediated stability of the sRNA-tmRNA complex, thereby increasing its inhibitory effect. Fluorescence measurements from the biosensor strains provided a quantitative assessment of *trans*-translation activity. The pAntisens_2 construct significantly reduced the fluorescence of biosensor 1, indicating successful inhibition of *trans*-translation. This reduction was not observed in biosensor 2, confirming the specificity of the inhibition. In contrast, the pAntisens_1 construct, lacking the MicC scaffold, showed a weaker, non-significative inhibitory effect, highlighting the importance of RNA stability in effective inhibition (Fig. 7).

**Figure 7:**
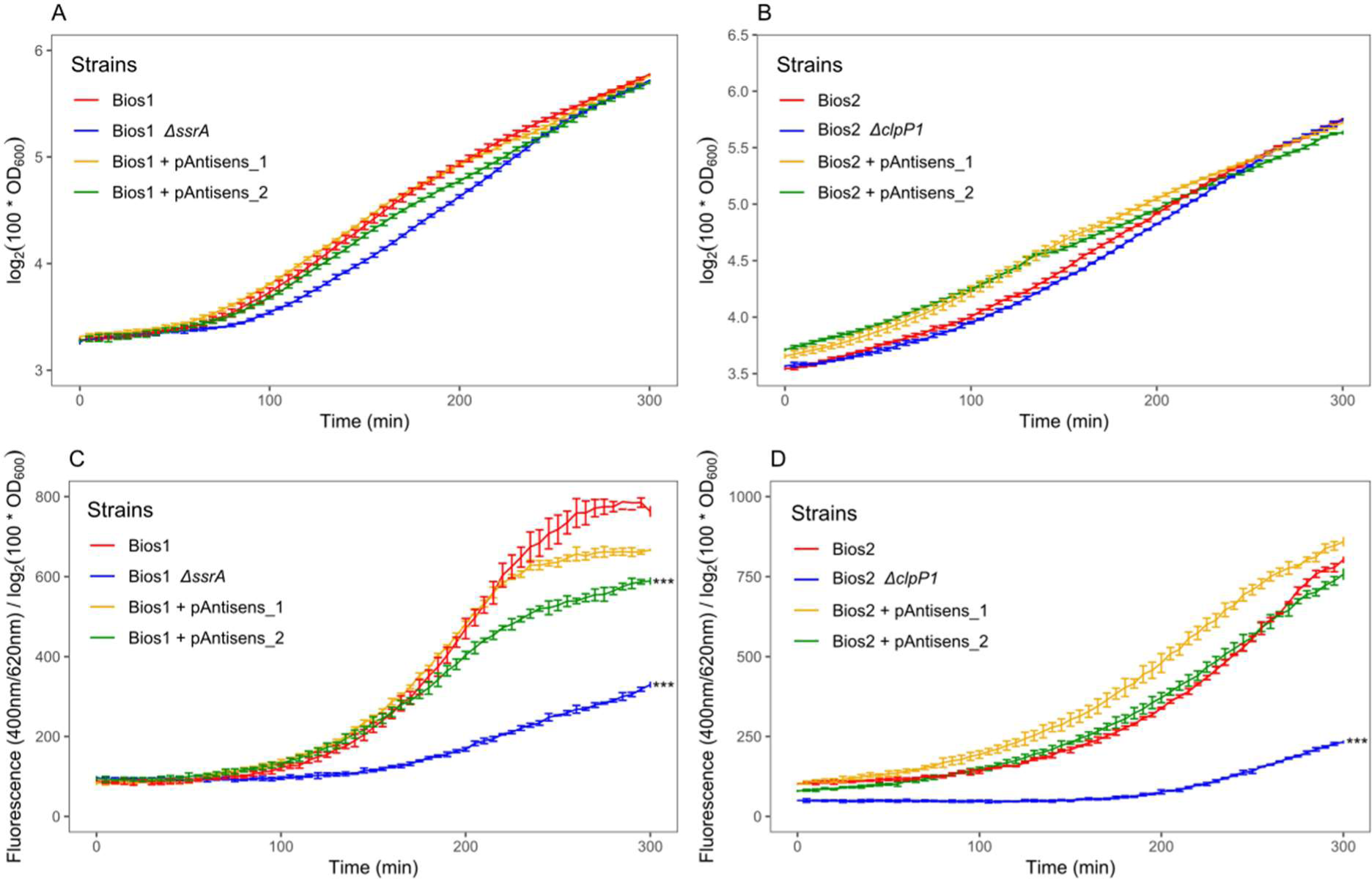
Effect of *trans-*translation inhibitors on the biosensors1 and biosensor2.Three repeats have been used for mean calculation and standard deviation. (A, B) Growth curve (optical density at 600nm according to time in minutes) of biosensor1 and biosensor2, respectively. (C, D) Fluorescence normalized by the optical density according to time in minutes of biosensor1 and biosensor2, respectively. (***) p-value <0.001 (Kruskal-Wallis test followed with Dunn test).

## 4 Discussion

We report the development of two biosensor strains capable of detecting novel anti-*trans*-translation compounds targeting *P. aeruginosa* PA14. These biosensors are based on the specific degradation of the ferrochelatase HemH, which converts the red fluorescent protoporphyrin IX into non-fluorescent heme. Biosensor 1 effectively identifies *trans*-translation inhibition, while biosensor 2 serves as a control by ensuring that observed effects are due to targeted inhibition rather than general cellular disruptions. This whole-cell-based system presents the advantage of targeting active molecules that are capable of penetrating cells. Additionally, it takes into consideration the capacity of efflux systems to expel these molecules, offering an advantage over an acellular system. The screening protocol has been miniaturized to 140μL in a microplate format, enabling automated kinetic monitoring of fluorescence. The rather small reaction volume and low cost highlights its potential for high-throughput screening of novel antimicrobial compounds.

Another advantage of this system’s working principle is that it can be applied to other bacteria in the ESKAPEE group, including *Staphylococcus aureus*, *Acinetobacter baumannii*, *Klebsiella pneumoniae*, and *Escherichia coli*. These bacteria have already demonstrated the ability to produce red fluorescence when exposed to violet light during fluorescence imaging (Jones et al., 2020). By using a genomic-based system rather than a plasmid-based one, the need for antibiotic resistance to maintain the system is eliminated. Furthermore, the *hemH*-based system not only removes the necessity for a dedicated insertion site but also leverages the bacteria’s intrinsic metabolic processes.

The next objective will be to select molecules that, at a concentration that does not significantly affect the growth of the biosensors, decrease the fluorescence of biosensor 1 without affecting the fluorescence of biosensor 2. This differential effect will help identify inhibitors that specifically target *trans*-translation rather than ClpP1 activity or general metabolic pathways. Overall, these biosensors offer a versatile and efficient tool for identifying novel antimicrobial agents, with broad applications across various bacterial species and the potential to accelerate the discovery of targeted therapies against drug-resistant pathogens.

## 5 Acknowledgments

This work was supported by grants RF20220503018/1/1/156 from the French Cystic Fibrosis Association “Vaincre la Mucoviscidose” and the french association “Gregory Lemarchal“; the Fondation pour la Recherche Médicale (Equipe FRM EQU202203014663) and Agence nationale de la Recherche ANR-23-CE18-0031-01.

