## supplementary data for "Developing Biosensors for Specific Assessment of *Trans*-translation in *Pseudomonas aeruginosa*"

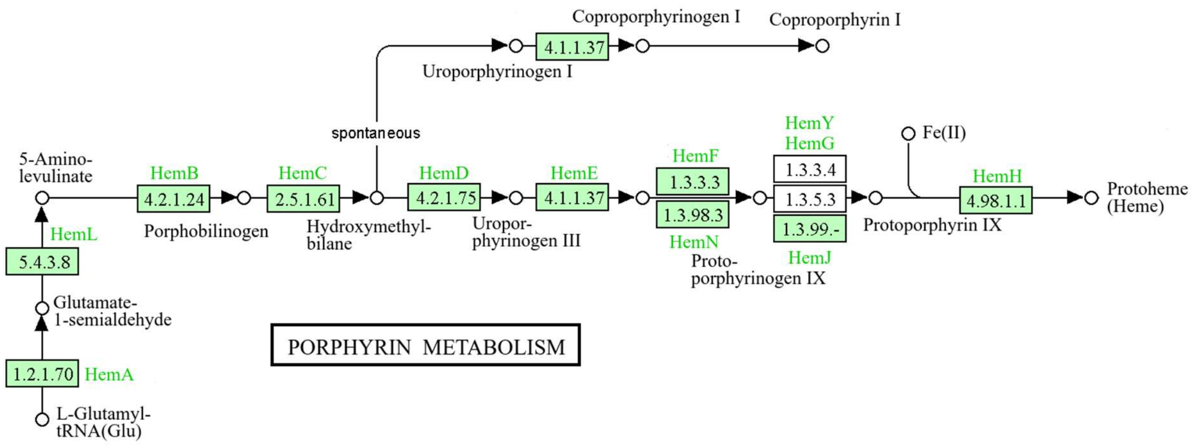


Figure S1: Porphyrin metabolism pathway of *P. aeruginosa* UCBPP-PA14 from L-Glutamyl-tRNA to Protoheme adapted from pathway map pau00860 (Kanehisa & Goto, 2000)

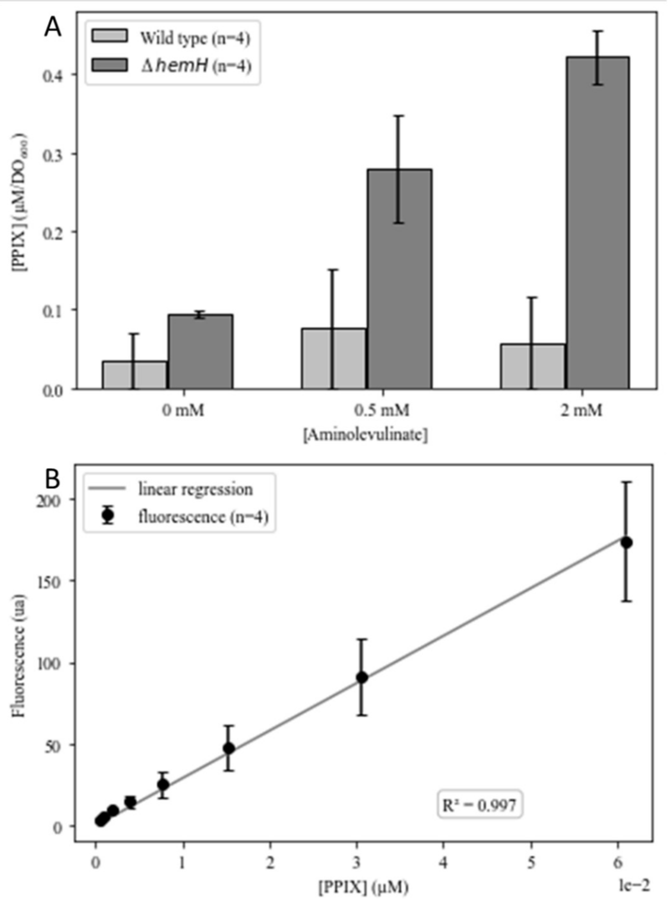


Figure S2: A) Concentration of PPIX, calculated with a standard curve (B) using fluorescence intensity of *P. aeruginosa* WT and *ΔhemH* lysate obtained from an exponential phase culture in CAA with 25μM hemin and with various concentrations of aminolevulinate.

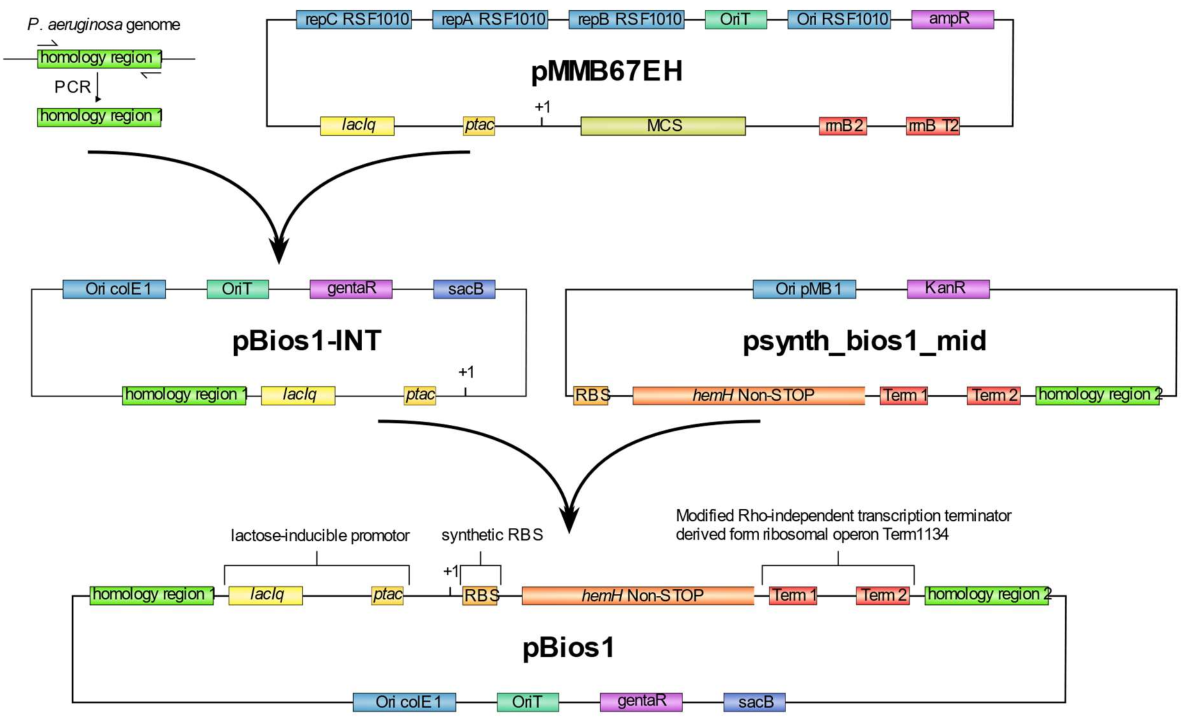


Fig S3: schematic representation of the construction of the plasmid pBios1.

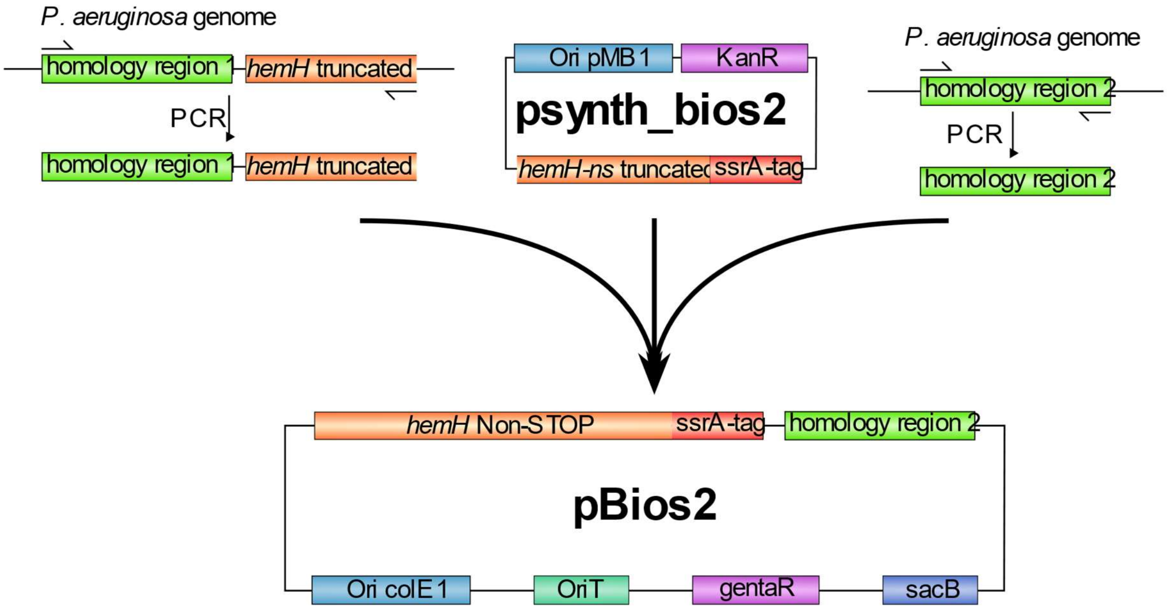


Fig S4: schematic representation of the construction of the plasmid pBios2.

Table S1: Synthetic sequences used to create the inhibitor plasmids.

| Name | DNA sequence<br>(antisense RNA) |
|---|---|
| Antisens_1 | GAATTCACGCGTTTATGACAACTTGACGGCTACATCATTCACTTTTTCTTCACAACCGGCACGGAACTCGCTCGGGCT<br>GGCCCCGGTGCATTTTTTAAATACCCGCGAGAAATAGAGTTGATCGTCAAAACCAACATTGCGACCGACGGTGGCGAT<br>AGGCATCCGGGTGGTGCTCAAAAGCAGCTTCGCCTGGCTGATACGTTGGTCCTCGCGCCAGCTTAAGACGCTAATCCC<br>TAACTGCTGGCGGAAAAGATGTGACAGACGCGACGGCGACAAGCAAACATGCTGTGCGACGCTGGCGATATCAAAATT<br>GCTGTCTGCCAGGTGATCGCTGATGTACTGACAAGCCTCGCGTACCCGATTATCCATCGGTGGATGGAGCGACTCGTT<br>AATCGCTTCCATGCGCCGCAGTAACAATTGCTCAAGCAGATTTATCGCCAGCAGCTCCGAATAGCGCCCTTCCCCTTG<br>CCCGGCGTTAATGATTTGCCCAAACAGGTCGCTGAAATGCGGCTGGTGCGCTTCATCCGGGCGAAAGAACCCCGTATT<br>GGCAAATATTGACGGCCAGTTAAGCCATTCATGCCAGTAGGCGCGCGGACGAAAGTAAACCCACTGGTGATACCATTC<br>GCGAGCCTCCGGATGACGACCGTAGTGATGAATCTCTCCTGGCGGGAACAGCAAAATATCACCCGGTCGGCAAACAAA<br>TTCTCGTCCCTGATTTTTCACCACCCCCTGACCGCGAATGGTGAGATTGAGAATATAACCTTTCATTCCCAGCGGTCG<br>GTCGATAAAAAAATCGAGATAACCGTTGGCCTCAATCGGCGTTAAACCCGCCACCAGATGGGCATTAAACGAGTATCC<br>CGGCAGCAGGGGATCATTTTGCGCTTCAGCCATACTTTTCATACTCCCGCCATTCAGAGAAGAAACCAATTGTCCATA<br>TTGCATCAGACATTGCCGTCACTGCGTCTTTTACTGGCTCTTCTCGCTAACCAAACCGGTAACCCCGCTTATTAAAAG<br>CATTCTGTAACAAAGCGGGACCAAAGCCATGACAAAATCGCGTAACAAAAGTGTCTATAATCACGGCAGAAAAGTCCA<br>CATTGATTATTTGCACGGCGTCACACTTTGCTATGCCATAGCATTTTTATCCATAAGATTAGCGGATCCTACCTGACG<br>CTTTTTATCGCAACTCTCTACTGTTTCTCCATATTAAGCAGCTAGAGCGTAGTTGTCGTCGTTGGCGTCGAC<br>(GACGCUUUUUAUCGCAACUCUCUACUGUUUCUCCAUAUUAAGCAGCUAGAGCGUAGUUGUCGUCGUUGGC) |
| Antisens_2 | ACGCGTTTATGACAACTTGACGGCTACATCATTCACTTTTTCTTCACAACCGGCACGGAACTCGCTCGGGCTGGCCCC<br>GGTGCATTTTTTAAATACCCGCGAGAAATAGAGTTGATCGTCAAAACCAACATTGCGACCGACGGTGGCGATAGGCAT<br>CCGGGTGGTGCTCAAAAGCAGCTTCGCCTGGCTGATACGTTGGTCCTCGCGCCAGCTTAAGACGCTAATCCCTAACTG<br>CTGGCGGAAAAGATGTGACAGACGCGACGGCGACAAGCAAACATGCTGTGCGACGCTGGCGATATCAAAATTGCTGTC<br>TGCCAGGTGATCGCTGATGTACTGACAAGCCTCGCGTACCCGATTATCCATCGGTGGATGGAGCGACTCGTTAATCGC<br>TTCCATGCGCCGCAGTAACAATTGCTCAAGCAGATTTATCGCCAGCAGCTCCGAATAGCGCCCTTCCCCTTGCCCGGC<br>GTTAATGATTTGCCCAAACAGGTCGCTGAAATGCGGCTGGTGCGCTTCATCCGGGCGAAAGAACCCCGTATTGGCAAA<br>TATTGACGGCCAGTTAAGCCATTCATGCCAGTAGGCGCGCGGACGAAAGTAAACCCACTGGTGATACCATTCGCGAGC<br>CTCCGGATGACGACCGTAGTGATGAATCTCTCCTGGCGGGAACAGCAAAATATCACCCGGTCGGCAAACAAATTCTCG<br>TCCCTGATTTTTCACCACCCCCTGACCGCGAATGGTGAGATTGAGAATATAACCTTTCATTCCCAGCGGTCGGTCGAT<br>AAAAAAATCGAGATAACCGTTGGCCTCAATCGGCGTTAAACCCGCCACCAGATGGGCATTAAACGAGTATCCCGGCAG<br>CAGGGGATCATTTTGCGCTTCAGCCATACTTTTCATACTCCCGCCATTCAGAGAAGAAACCAATTGTCCATATTGCAT<br>CAGACATTGCCGTCACTGCGTCTTTTACTGGCTCTTCTCGCTAACCAAACCGGTAACCCCGCTTATTAAAAGCATTCT<br>GTAACAAAGCGGGACCAAAGCCATGACAAAATCGCGTAACAAAAGTGTCTATAATCACGGCAGAAAAGTCCACATTGA<br>TTATTTGCACGGCGTCACACTTTGCTATGCCATAGCATTTTTATCCATAAGATTAGCGGATCCTACCTGACGCTTTTT<br>ATCGCAACTCTCTACTGTTTCTCCATATTAAGCAGCTAGAGCGTAGTTGTCGTCGTTGGCTTTCTGTTGGGCCATTGC<br>ATTGCCACTGATTTTCCAACATATAAAAAGACAAGCCCGAACAGTCGTCCGGGCTTTTTTTTT<br><br>(AGCAGCUAGAGCGUAGUUGUCGUCGUUGGCUUUCUGUUGGGCCAUUGCAUUGCCACUGAUUUUCCAACAUAUAAAAA<br>GACAAGCCCGAACAGUCGUCCGGGCUUUUUUUUU) |
